# Systematic comparison of estimates of transcription factor activity by ATAC-seq and multiplexed reporter assays

**DOI:** 10.64898/2026.01.05.697632

**Authors:** Max Trauernicht, Vinícius H. Franceschini-Santos, Teodora Filipovska, Bas van Steensel

## Abstract

Transcription factors (TFs) are central to gene regulation and play critical roles in development, cellular homeostasis and disease. The ability to accurately measure TF activity is essential to understanding how TFs respond to signals and regulate target genes. In one commonly used approach, activities of TFs are computationally inferred from genome-wide chromatin accessibility data (ATAC-seq). However, it has remained unclear how well these inferences reflect actual regulatory activity of TFs. An alternative approach employs a collection of synthetic reporters that are designed to each probe the regulatory activity of a single TF. In this study, we systematically compared TF activities as inferred by ATAC-seq with those measured by multiplexed reporters, across diverse perturbations known to alter specific TF activities. We observed considerable overlap between the two methods, but also notable discrepancies. Our findings suggest that reporter assays and chromatin-based inference capture distinct aspects of TF function: reporter assays are more sensitive to signal-responsive TFs, while ATAC-seq better detects chromatin-modifying TFs.

## INTRODUCTION

Transcription factors (TFs) are fundamental regulators of gene expression. They determine cell identity, coordinate developmental programs, and play critical roles in the onset and progression of numerous diseases. As such, understanding TF activity is key to decoding cellular behavior and disease mechanisms. Ideally, we would like to measure the activity of all TFs in parallel within a single cell to gain a comprehensive view of transcriptional regulation. However, this is a considerable challenge. TFs exert their effects by binding to specific sites in promoters and enhancers, often within densely packed regulatory regions where dozens of different TFs can act simultaneously. These complex and combinatorial interactions make it difficult to isolate and interpret the functional contribution of individual TFs. Therefore, to truly understand TF function within the genome, we require experimental methods and computational strategies capable of deconvoluting these intricate regulatory networks.

While the abundance of a TF can provide some indication of its potential activity within a cell and can be approximated by measuring the expression level of the mRNA that encodes the TF (1), expression levels alone often fail to accurately reflect a TF’s true functional activity. This is because mRNA abundance does not always correlate with protein abundance, and many TFs are further regulated by upstream signaling pathways and post-translational modifications that modulate their activity. For example, nuclear receptors are activated through binding of specific ligands (2), CREB1 (cAMP responsive element binding protein 1) becomes transcriptionally active through cAMP-mediated phosphorylation (3), and HSF1 (heat shock transcription factor 1) is activated by trimerization in response to proteotoxic stress (4). These activation mechanisms highlight that identical TF mRNA or protein levels can correspond to vastly different transcriptional outcomes depending on the presence or absence of activating signals.

An alternative to measuring TF abundance is to infer TF activity from their DNA-binding patterns. While direct TF binding assays like ChIP-seq (chromatin immunoprecipitation sequencing) (5) provide insight, they are labor-intensive and limited to one single TF at a time. Instead, chromatin accessibility assays such as ATAC-seq (assay for transposase-accessible chromatin with sequencing) (6) are increasingly used to infer TF activity. This is based on the assumption that TFs can open up the chromatin locally, often by displacing one or more nucleosomes (7–11). Overall activity of a TF is thus inferred by scoring the genome-wide correlation between the presence of its binding motif and the occurrence of open chromatin. This correlation analysis can be repeated for all TF binding motifs. ATAC-seq is simple, requires little input material, and can be applied to single cells, making it an attractive method. Several computational tools now exist to infer TF activity from ATAC-seq data (12–15), and these are widely used across biological disciplines (16–21). These tools enable efficient and scalable computation of activity scores for any given TF.

However, this approach provides an indirect measure of TF function. ATAC-seq captures the ability of a TF to bind DNA and alter chromatin accessibility, not necessarily its transcriptional output. This introduces biases: for example, pioneer TFs, which can bind and open closed chromatin, are more likely to be captured by ATAC-seq. In contrast, TFs that require pre-existing open chromatin may be underrepresented. Additionally, some TFs may bind accessible regions not because they actively shape chromatin, but simply because the chromatin was opened by other factors. As a result, TFs that associate with constitutively open regions, such as housekeeping promoters, may seem highly active despite having little direct transcriptional influence.

These limitations raise a central question: what does ATAC-seq truly measure about TFs? And what should we aim to measure to accurately capture their functional roles? Ideally, we would measure transcriptional activity of TFs directly, isolating the specific contribution of each TF without interference from combinatorial binding or chromatin context. Recent advances in reporter technology now make this possible. Massively parallel reporter assays (MPRAs) have been developed to measure the transcriptional activity of many elements in parallel, using barcoded reporters and sequencing-based readout (22). We previously generated highly optimized “prime” TF reporters (23), enabling the parallel quantification of the transcriptional activity of 100 TFs in a simple assay (24).

In this study, we systematically compared TF activity inferred from ATAC-seq and TF activity measured directly using prime TF reporters. By comparing these two approaches, we assessed their respective strengths and limitations, and clarified what each method captures about TF function. To do so, we applied various single-factor perturbations to cells, as well as induction of differentiation, and then compared both methods in identifying differential TF activities. We found that prime reporters were generally more sensitive to changes in TF activity and were particularly effective in detecting signal-responsive TFs. In contrast, ATAC-seq preferentially detected chromatin-modifying TFs, such as pioneer factors. Together, these insights help refine our understanding of how TF activity is best measured, and inform future strategies for studying transcriptional regulation.

## RESULTS

### Two methods to measure TF activity: ATAC-seq-based inference and prime TF reporters

To systematically evaluate the ability to quantify TF activity, we performed side-by-side comparisons of a multiplexed TF reporter assay and chromatin accessibility profiling by ATAC-seq (**Figure 1a**). For the reporter-based approach, we employed the previously established “prime” TF reporter library (24), which detects the transcriptional activity of 100 TFs using optimized barcoded TF reporters. Notably, the prime TF reporter library contains 62 TFs that are classified as “prime” TF reporters (i.e., reporters with strong evidence for TF-specificity) and 38 TFs classified as non-prime. To ensure broad and unbiased coverage, we included all 100 reporters in our analysis. For simplicity, we refer to this full set as the prime TF reporter library, despite the inclusion of non-prime TFs. In the prime TF reporter assay, the TF reporter plasmid library is transfected into cells, and transcription from these reporters is measured by quantifying the barcodes in the mRNA using high-throughput sequencing. Reporter-based TF activities were then computed using *primetime* (24), a tool to calculate differential TF activities from barcode counts using the prime TF reporter library. For chromatin accessibility-based TF activity detection, we performed the standard ATAC-seq protocol and quantified TF activity using chromVAR (12), a tool that measures TF-associated accessibility (**Figure 1a**). We selected chromVAR with minor modifications for quantification because it was previously shown to outperform alternative approaches in capturing perturbation-induced changes in TF activity (25).

**Figure 1:**
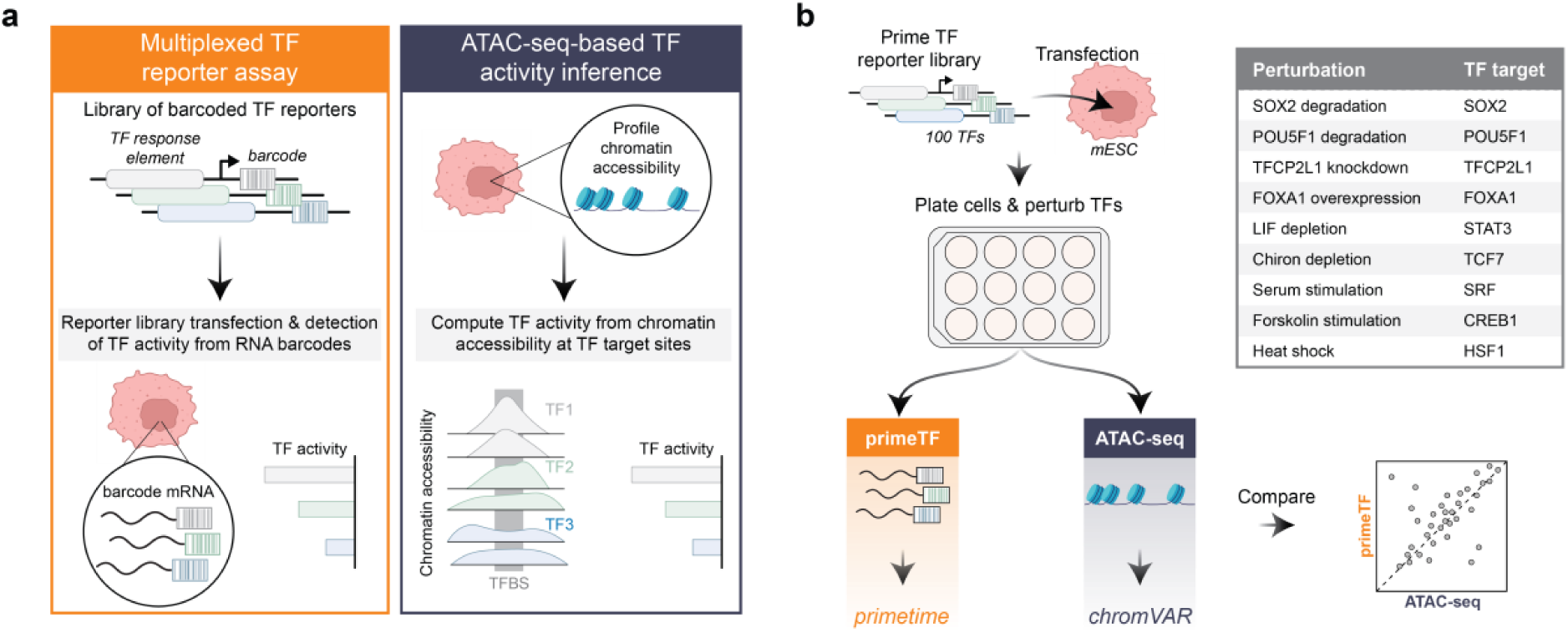
Systematic comparison of TF activity computation by ATAC-seq and TF reporter activities. a) Scheme of the two different approaches to measure TF activity – multiplexed TF reporter assay (left panel) and ATAC-seq-based TF activity inference (right panel). TFBS = TF binding site. b) Scheme of the experimental layout to compare multiplexed TF reporter assay and ATAC-seq-based TF activity inference. mESCs are transfected with the prime TF reporter library, plated into wells and perturbed using a variety of perturbation conditions, after which ATAC-seq and TF reporter barcode sequencing is performed. TF activities are then computed from ATAC-seq using *chromVAR*, and from TF reporter barcode counts using *primetime*, and then compared to each other.

### Systematic TF perturbations to benchmark TF activity detection methods

To ensure a fair comparison between the prime TF reporter assay and ATAC-seq, all measurements were performed in the same cellular background following transfection of the reporter library, thereby controlling for transcriptional changes that may be induced by transfection-related stress. To benchmark assay sensitivity and specificity, we transfected the reporter library into mouse embryonic stem cells (mESCs) and then systematically perturbed nine TFs (**Figure 1b**). These TFs were selected based on their presence as targets in the reporter library. Perturbations included the degradation of core pluripotency factors SOX2 and POU5F1 (both 24 hours) using degron-tagged cell lines (26), knockdown of TFCP2L1 (24 hours), and overexpression of FOXA1 (48 hours). In addition, we employed signaling-based perturbations to modulate STAT3 and TCF7 activity via withdrawal of LIF and Chiron (both 24 hours), respectively. Finally, we stimulated SRF (serum response factor), CREB1, and HSF1 through serum exposure (6 hours), treatment with cAMP-inducing forskolin (6 hours), and heat shock (3 hours), respectively. Following these perturbations, mESCs were harvested in parallel for either ATAC-seq, or RNA extraction and barcode sequencing for reporter activity profiling. TF activities were then computed using *primetime* and *chromVAR* following sequencing. For fair comparison of the two techniques, we use the exact same TF binding motifs for the ATAC-seq analysis as the ones used as binding sites in the TF reporters. This experimental design enabled direct comparison of TF activity readouts between assays, providing a framework for evaluating their respective sensitivity and biological relevance.

### chromVAR detects changes in chromatin accessibility around TF binding sites (TFBSs)

We first validated our experimental setup by quantifying accessibility of SOX2 binding sites in the SOX2 degradation condition. SOX2 is a known pioneer TF (26–28), and prior research demonstrated that degradation of SOX2 results in rapid reduction of chromatin accessibility at genomic SOX2 binding sites (26). Given its established role, SOX2 serves as an ideal benchmark for validating ATAC-seq-based TF activity detection. Consistent with previous findings, we detected substantially reduced chromatin accessibility at genomic SOX2 binding sites following SOX2 degradation (**Figure 2a**). We then employed chromVAR to calculate bias-corrected deviation scores for all 100 TF motifs that are present in the prime TF reporter library. These deviation scores quantify the degree to which observed accessibility surrounding each motif deviates from expected accessibility across all cells, with technical biases such as GC content appropriately controlled for. For statistical comparison between conditions, we used the limma framework (29) to assess differential deviation scores. This method was previously shown to be a robust method for the differential analysis of chromVAR-computed deviation scores (25). As anticipated, SOX2 binding sites exhibited significantly reduced chromVAR deviation scores following SOX2 degradation (**Figure 2b**). We validated our computed deviation scores through multiple quality controls. First, differential deviation scores between replicates following SOX2 degradation showed strong correlations (*R* = 0.92-0.97, **Figure EV1a, b**). Moreover, cross-validation with previously published ATAC-seq data (26) using identical mESC SOX2 degron cells revealed high correlation (*R* = 0.91, **Figure EV1c**). Additionally, the computed differential chromVAR deviation scores correlated well with differential log odds ratios, a more basic metric of motif accessibility (*R* = 0.79, **Figure EV1d**). Together, these results confirm both our experimental data quality and the robustness of chromVAR in the detection of differential TF activities.

**Figure 2:**
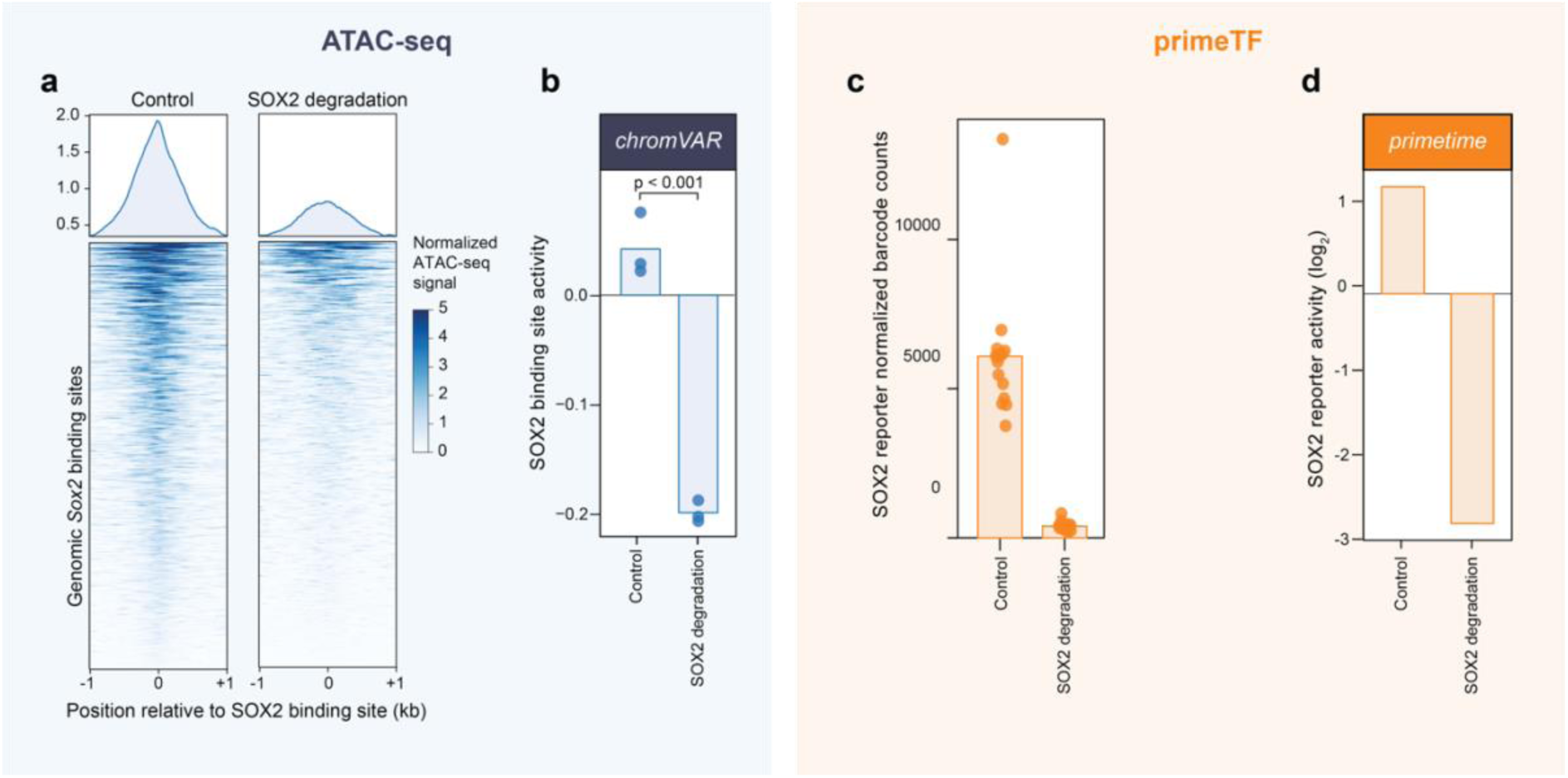
TF activity detection by *chromVAR* and *primetime*. a) Normalized ATAC-seq signal 2 kb around the 1000 best-matching SOX2 binding sites in the genome (see Methods) in control cells (left panel) and upon SOX2 degradation (right panel). Upper plot shows average. Displayed is the average across three biological replicates. b) Bias-corrected deviation scores of the SOX2 binding site computed by chromVAR in control cells and upon SOX2 degradation. Displayed are the three biological replicates (dots) and the mean across those (bar). Statistical significance between conditions was tested using limma. c) Barcode counts (normalized by sequencing depth) of the SOX2 reporters in control cells and upon SOX2 degradation. Displayed are the counts of the five barcodes of the SOX2 reporter and the three biological replicates (15 dots) and the mean across those (bar). d) Reporter activities and significance between the two conditions as computed by *primetime*.

### primetime *detects changes in TF activity upon perturbation*

We subsequently quantified changes in SOX2 reporter activity following SOX2 degradation using the prime TF reporter library. We found a substantial reduction (approximately 15-fold) in barcode counts for SOX2 reporters upon SOX2 degradation (**Figure 2c**). We then applied *primetime* to compute TF activity scores and perform differential analysis to statistically evaluate the effect of the perturbation. This confirmed a significant reduction in SOX2 reporter activity following degradation (**Figure 2d**). As part of the standard quality control steps of the *primetime* workflow, we further confirmed low technical noise between measured barcodes (**Figure EV1e**) and high biological replicate reproducibility (**Figure EV1f**). These results demonstrate the robustness of the assay and the effectiveness of *primetime* in capturing TF activity changes in response to targeted perturbations.

### TF reporters demonstrate higher sensitivity in detecting perturbations of target TFs

After validating the sensitivity of both assays to detect changes in TF activity upon perturbation, we quantified the activity changes of the targeted TFs across all nine perturbation conditions using both ATAC-seq and primeTF. (**Figure 3a**). primeTF detected significant and directionally accurate changes in all nine cases, with fold-changes ranging from 2.6-fold (SRF upregulation upon serum stimulation) to 63-fold (HSF1 upregulation upon heat shock). In contrast, the ATAC-seq-based readout detected significant activity changes in seven of the nine targeted TFs, failing to capture SRF upregulation in response to serum stimulation and CREB1 upregulation following forskolin treatment. The most substantial ATAC-seq activity difference of target TFs were observed upon degradation of POU5F1 and SOX2. To further compare assay performance, we ranked TFs by their perturbation response across all 100 investigated TFs (**Figure 3b**). In six of the nine perturbations - SOX2 degradation, FOXA1 overexpression, LIF depletion, TFCP2L1 knockdown, Chiron depletion, and POU5F1 degradation - both ATAC-seq and primeTF identified the targeted TF as the top or second most responsive. However, in the heat shock, forskolin, and serum stimulation conditions, ATAC-seq failed to prioritize the expected primary targets, ranking them only 4th, 14th, and 85th, respectively.

**Figure 3:**
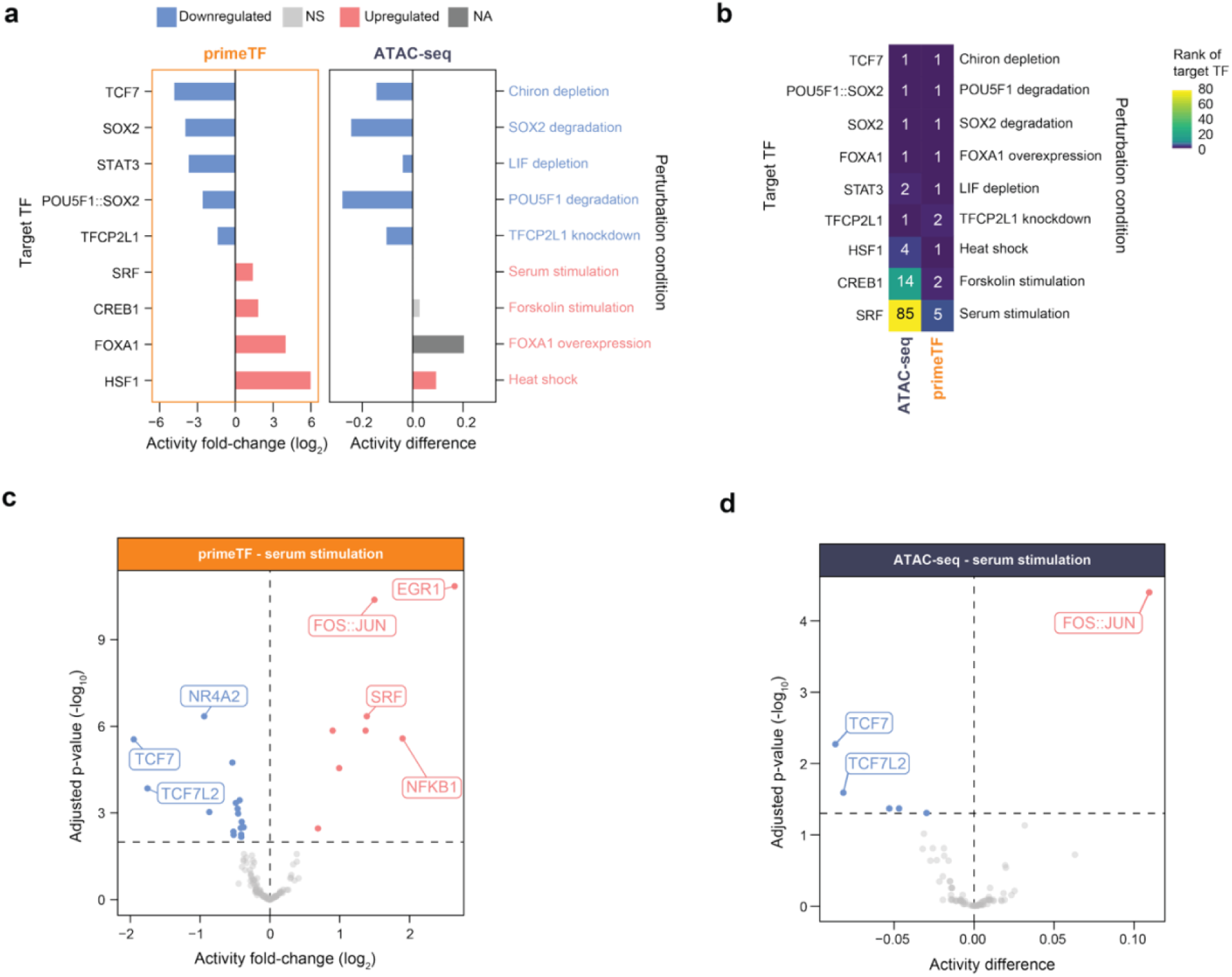
primeTF detects target TF activity changes with higher sensitivity. a) Fold-change (log_2_) in *primetime* reporter activity (left panel) and difference in *chromVAR* deviation score (right panel) of the nine perturbed target TFs. Significance of FOXA1 deviation score differences could not be tested since only one replicate was available. b) Ranks of the target TFs sorted by the perturbation effect as calculated by the adjusted p-value (-log_10_) multiplied by the activity fold-change or deviation score difference. c) Vulcano plot of the changes in reporter activity following serum stimulation as calculated by *primetime*. d) Same as c) showing the differences in *chromVAR* deviation scores.

### Both methods pick up broader perturbation responses

To gain deeper insight into what each method captures, and to reveal potential differences in sensitivity and bias, we quantified activity changes for all 100 TFs in parallel across each perturbation, rather than limiting our analysis to the targeted TF alone. As an illustrative example, serum stimulation elicited a widespread transcriptional response, with primeTF identifying 8 TFs as upregulated and 16 as downregulated (**Figure 3c**), and ATAC-seq detecting 1 upregulated and 5 downregulated TFs (**Figure 3d**). primeTF identified next to primary target TF SRF also other TFs that are known downstream targets of serum response (e.g. EGR1, FOS::JUN (30,31)). ATAC-seq, on the other hand, did not identify an upregulation of primary target SRF, but only of its downstream target FOS::JUN. Both methodologies consistently detected significant downregulation of TCF7/TCF7L2 following serum stimulation, demonstrating concordance in specific aspects of the cellular response.

### Comparing TF reporter assay and ATAC-seq across 100 TFs and nine TF perturbations conditions

To systematically evaluate the performance of the two assays, we analyzed the activity changes of all 100 measured TFs across the nine distinct perturbation conditions (see **Dataset EV1**). For an unbiased comparison, we constructed comprehensive correlation plots for each perturbation, enabling side-by-side identification of the most responsive TFs by both methods (**Figure 4a**). In several cases, both methods concordantly identified the perturbed TF as the most strongly affected, often without widespread secondary effects. For example, Chiron depletion, TFCP2L1 knockdown, and FOXA1 overexpression led to pronounced activity changes in their respective target TFs, with minimal effects on unrelated TFs. However, for some perturbations, as mentioned before for serum stimulation (**Figure 3c, d**), changes in TF activity disagreed between the two methods. For instance, upon LIF depletion, primeTF detected a very strong downregulation of only target TF STAT3 without affecting any other TFs, while ATAC-seq detected moderate downregulations of many TFs, including STAT3. Another example is SOX2 and POU5F1 degradation, upon which both methods detected downregulation of SOX2 and POU5F1::SOX2, respectively, but reported largely disagreeing secondary changes in TF activity. Notably, ATAC-seq revealed a marked decrease in POU5F1 levels following its degradation, while the activity of the POU5F1 primeTF reporter showed only a slight reduction. This suggests that the POU5F1 reporter is less sensitive to changes in POU5F1 levels compared to the POU5F1::SOX2 reporter. What is more, primeTF identified CREB1 and HSF1 as the most upregulated TFs following forskolin stimulation and heat shock, respectively, whereas ATAC-seq failed to detect any changes in response to forskolin and did not rank HSF1 among the top responders to heat shock. To exclude that these discrepancies are due to the poor performance of the included non-prime reporters in the primeTF reporter library, we generated the same correlation plots without including non-prime reporters, which yielded highly similar correlation plots for all conditions (**Figure EV2a**). Collectively, these findings indicate that the primeTF reporter assay outperforms ATAC-seq in accurately identifying the primary TF target across most perturbations. Beyond these major differences, both methods also report numerous distinct secondary changes, the interpretation of which may depend on differences in sensitivity, specificity, and the underlying biases of each approach.

**Figure 4:**
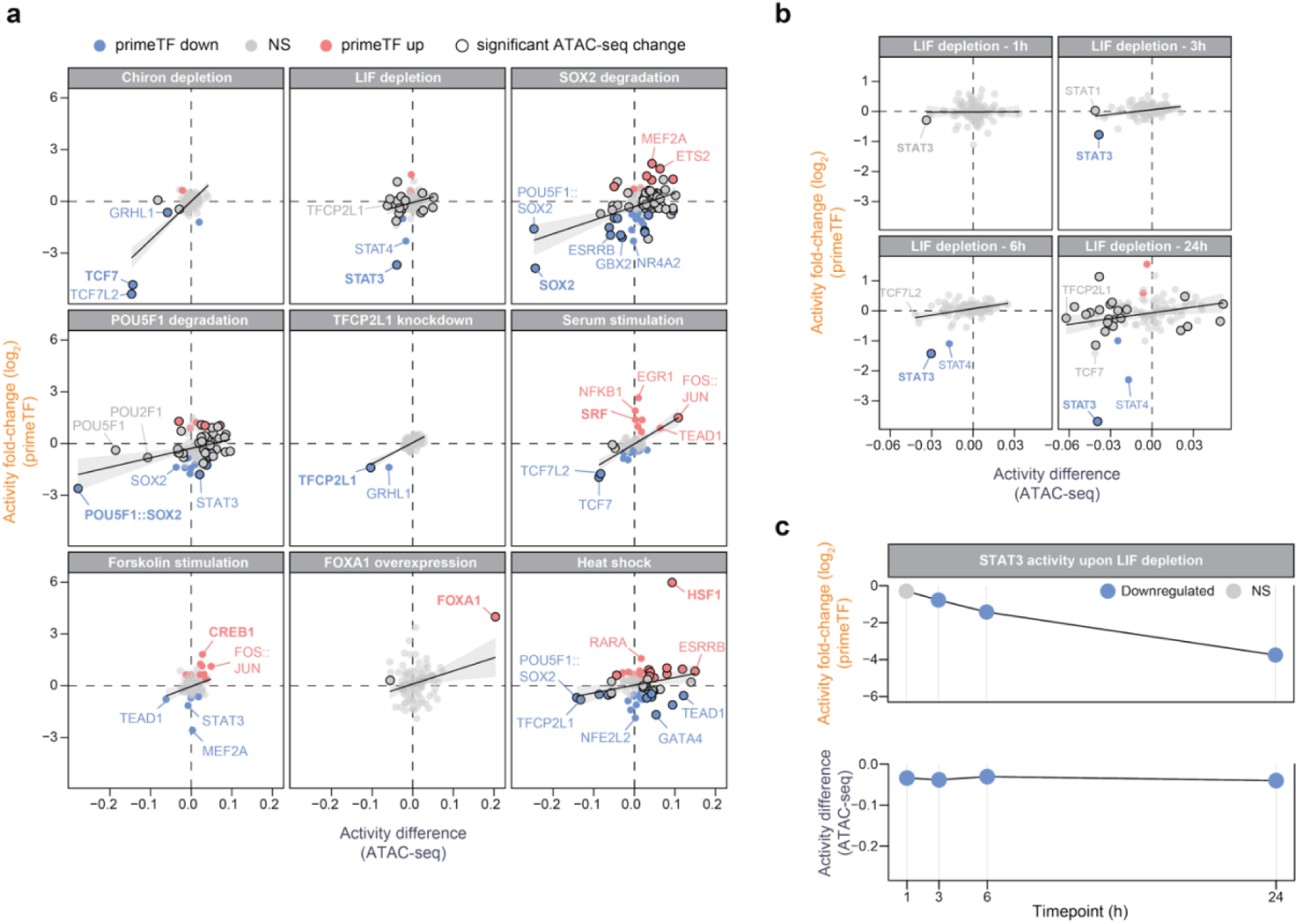
Changes in secondary TFs differ between ATAC-seq and primeTF. a) Correlation between difference in *chromVAR* deviation score and fold-change (log_2_) in primetime activity across the nine perturbation conditions. Point fill color indicates statistical significance based on *primetime*, stroke (outline) denotes significance according to *chromVAR*. b) Same as a) but for different time-points after LIF depletion. c) Changes in STAT3 activity at different time-points after LIF depletion measured by *primetime* and *chromVAR*. Y-axis is scaled according to maximum effect sizes observed across all perturbations per assay.

### Discrepancies may be explained by different time resolutions

One potential explanation for the observed discrepancies between the TF reporter assay and ATAC-seq is the differing temporal resolution inherent to each method. While ATAC-seq captures a snapshot of chromatin accessibility at the time of cell harvest, primeTF reflects TF activity over a defined window of time, shaped by both transcriptional induction and mRNA degradation. This results in a semi-integrated readout that balances dynamic responsiveness with signal accumulation. To directly assess the impact of this difference, we performed a time-course experiment using LIF depletion as a perturbation of STAT3 activity. We collected samples at 1, 3, 6, and 24 hours post-LIF withdrawal and profiled them using both ATAC-seq and primeTF. As early as 1 hour post-LIF removal, ATAC-seq reported a measurable decrease in STAT3 activity, which reached its maximum downregulation by 3 hours and remained stable through 24 hours (**Figure 4b, c**). Importantly, significantly changed activities of TFs other than STAT3 were only detected after 24 hours. In contrast, the TF reporter assay did not detect STAT3 downregulation at 1 hour but began to show reduced activity at 3 hours, with a continued progressive decrease up to 24 hours. These results underscore that ATAC-seq provides a near-instantaneous measure of chromatin changes, while primeTF reflects a time-integrated transcriptional activity. However, this temporal distinction alone does not fully explain the differences in sensitivity between the two methods. Notably, at both 6 and 24 hours, primeTF identified STAT3 as the most significantly downregulated TF with a large effect size, whereas ATAC-seq reported comparable effect sizes for several other TFs, failing to clearly prioritize STAT3. This suggests that, in addition to capturing temporal dynamics differently, primeTF may provide superior sensitivity in identifying primary TF targets across extended perturbation windows.

### Both methods detect loss of pluripotency TF activity upon differentiation

While benchmarking ATAC-seq and primeTF using controlled single-factor perturbations offered clear insights into the behavior of each method, applying them in a more complex setting, such as differentiation into another cell type, allows to explore their performance in a biologically relevant, but less defined, context. Differentiation typically involves coordinated changes in the activity of multiple TFs, making it a relevant context for evaluating these tools. To assess this, we differentiated mESCs into mouse neural precursor cells (mNPCs) (32) and performed both ATAC-seq and the prime TF reporter assay in these two cell types. As before, we then computed changes in TF activity between mESCs and mNPCs using both *chromVAR* and *primetime* (**Figure 5a**). We first checked if the changes picked up by ATAC-seq using the 100 TFBSs included in this study were capturing a wide range of changes compared to all known human TFBSs. We found that the 100 TFBSs were significantly enriched for high deviation scores relative to the rest of the TF motif database (**Figure EV2b**). While still missing biologically relevant changes, this indicates that our selected set of 100 TFs captures a large proportion of the chromatin accessibility changes occurring during differentiation. We then focused our analysis on key pluripotency TFs, which are expected to be active in mESCs but downregulated upon differentiation into mNPCs. primeTF successfully identified the downregulation of all pluripotency TFs upon differentiation. In contrast, ATAC-seq detected downregulation of most pluripotency TFs, but failed to identify the downregulation of key pluripotency TFs STAT3 and POU5F1 (**Figure 5b**).

**Figure 5:**
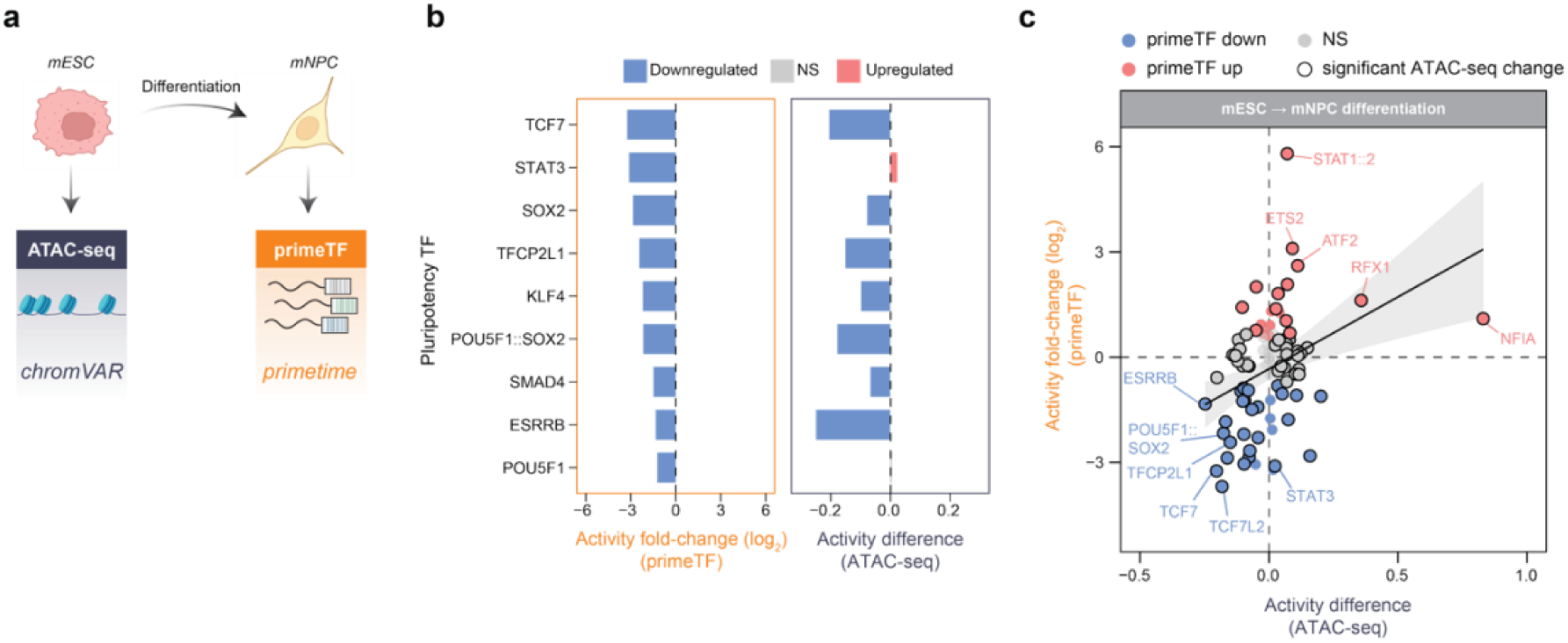
Differential TF activity detection upon differentiation. a) Scheme of experimental workflow. mESCs were differentiated into mNPCs. As before, ATAC-seq and TF reporter barcode sequencing were performed in both cell types to compute and compare TF activities from *chromVAR* and *primetime*. b) Changes in activity of known pluripotency TFs upon mNPC differentiation measured by *primetime* (left panel) and *chromVAR* (right panel). c) Correlation between changes in *chromVAR* deviation score and *primetime* activity upon mESC differentiation into mNPCs.

### Differences between ATAC-seq and primeTF

*both methods are biased towards certain TFs*. We then analyzed the activity differences of all 100 TFs. We found that the changes in activity detected by ATAC-seq and primeTF correlate moderately (rho = 0.36), with many TFs changing activity significantly in the same direction (**Figure 5c**). However, there were notable differences between the methods for specific TFs. For example, STAT1::2 was upregulated 56-fold according to primeTF, but only showed a slight increase in activity with ATAC-seq. STAT1::2 responds to interferon pathways, which are repressed in mESCs to maintain pluripotency and avoid differentiation signals, and becomes upregulated in differentiated cells such as mNPCs (33). This may explain the elevated activity detected with the TF reporter assay. ETS2 and ATF2 exhibit a similar pattern, with a strong upregulation in primeTF but only a modest increase in ATAC-seq. In contrast, ATAC-seq detected a strong upregulation of NFIA in mNPCs (three times greater than the change induced by SOX2 degradation on SOX2 activity) while primeTF detected only a two-fold NFIA upregulation. NFIA plays a key role in neural and glial differentiation (34) and is highly expressed in mNPCs but not mESCs. It is likely that NFIA acts as a pioneer TF, in line with previous findings (35), inducing substantial chromatin remodeling upon expression, but may function as a weaker transcriptional activator on its own. We note that NFIA is not classified as a “prime” TF reporter, which might explain the relatively poor performance of the reporter. Focusing solely on the 62 prime reporters - excluding the NFIA outlier - resulted in a similar correlation between the two methods (rho = 0.37, **Figure EV2c**). Together, the differences detected upon differentiation by each method underscore the distinct strengths and weaknesses of each method, revealing that each tool tends to highlight specific sets of TFs with unique properties.

## DISCUSSION

Genome-wide accessibility profiles generated by ATAC-seq are now widely used to infer TF activity (36–40), dissect cis-regulatory logic (41), and reconstruct gene regulatory networks (42,43). However, it is essential to recognize that chromatin accessibility is an indirect proxy. While useful, it predominantly reflects the chromatin-opening potential of TFs and may overlook other critical modes of regulation. To clarify what ATAC-seq truly captures and to better define "TF activity", we systematically benchmarked ATAC-seq-derived activity against direct transcriptional activity measured by a multiplexed TF reporter assay. We found substantial agreement between the two methods but also revealed clear biases and discrepancies that illuminate the unique strengths of each approach.

A key finding was the enhanced sensitivity of the TF reporter assay in detecting signal-responsive TFs. Upon various perturbations, including heat shock, cAMP induction, serum stimulation, and LIF depletion, primeTF detected strong changes in activity of HSF1, CREB1, SRF, and STAT3, respectively. These TFs are canonical early responders whose activation is typically driven by post-translational modifications rather than changes in abundance or chromatin remodeling (30,31,44–46). ATAC-seq failed to identify these TFs as primary target upon their perturbation. Since many of these TFs bind to already accessible chromatin, their activity may not manifest as shifts in accessibility detectable by ATAC-seq. This limitation of chromatin-based assays aligns with prior findings that numerous genes are transcriptionally activated without accompanying changes in nearby chromatin accessibility (47). Our results reinforce that reporter-based methods provide a valuable complement for capturing dynamic, signal-dependent transcriptional responses.

Conversely, ATAC-seq demonstrated strong detection upon FOXA1, TCF7, SOX2, and POU5F1 perturbation, and prioritized NFIA upon mNPC differentiation in contrast to the TF reporter assay. These TFs are classic pioneer TFs (26,48–52), known to open compact chromatin, leaving clear accessibility footprints that ATAC-seq can detect. In line with this, ATAC-seq picked up pioneer TF FOS::JUN (53,54) upon serum stimulation, while failing to detect signal-responsive TFs like SRF. Hence, our results indicate that ATAC-seq based detection is biased towards detecting chromatin-modifying pioneer TFs. The TF reporter assay, on the other hand, appeared to underreport some of these TFs, such as NFIA upon mNPC differentiation. This is likely due to limitations in the reporter design or weak transcriptional activity in isolation. Interestingly, the TF reporters still captured substantial activity changes for several pioneer TFs, suggesting that it does not completely overlook these TFs, contrary to previous reports (55,56).

Through our LIF depletion time-course we found that the time resolution of the two assays is different: ATAC-seq picks up changes instantaneously, whereas the TF reporters measure responses with a delay. Thus, in some of our perturbation experiments ATAC-seq might have already ‘missed’ the activation of a target TF because it is an early response TF that acts quickly upon activation and then returns to normal levels (e.g., SRF (57)). Confirming this, we often find changed ATAC-seq signals of known downstream targets of the perturbed TFs. For instance, upon Chiron depletion, ATAC-seq detected changed activity of TFCP2L1, which is a known downstream target of TCF7 (58). Similarly, ATAC-seq identified TEAD1 involved in the heat shock response, as previously reported (59). However, our time-course experiment also revealed that early responsive TFs are not completely missed by ATAC-seq. Rather, ATAC-seq has a tendency to underestimate their importance, likely because they are not substantially altering chromatin accessibility. To obtain a more immediate readout of TF reporter activity, nascent transcription profiling, such as 4sU labeling (60–62), could be used. However, this approach may be technically challenging.

In this study, we primarily focused on single-factor perturbations targeting transcription factors that are part of the “prime” set in the reporter library. These TFs were deliberately chosen because their corresponding reporters had previously been validated to respond to perturbation. As such, our analysis serves more as a benchmark to evaluate whether ATAC-seq can match the performance of these well-characterized TF reporters, rather than a test of the reporter system itself. To assess the relative strengths and limitations of both methods more comprehensively, future studies should also include perturbations of TFs that fall outside the “prime” category. Such an approach would offer a more unbiased comparison of the two techniques in detecting TF activity changes. We note that the differentiation experiment included in this study serves as an example of such an unbiased setting. However, this system is inherently more complex and lacks a clearly defined ground truth for expected TF activity changes, making direct benchmarking more challenging.

Although our study focused on chromVAR, numerous other tools infer TF activity from ATAC-seq data. However, chromVAR has recently been shown to outperform many of these methods (25), suggesting that similar or greater limitations may exist elsewhere. Another consideration is that chromVAR is mostly used with single-cell ATAC-seq data. The drawbacks identified here also apply to these analyses. However, it is not yet feasible to do single-cell TF reporter measurements. Developing methods to measure multiplexed TF activity at single-cell resolution would be a major advancement, enabling researchers to directly link signaling responses and transcriptional output at the level of individual cells.

In summary, we conclude that the two methods offer distinct and complementary perspectives on TF function. ATAC-seq excels at identifying TFs that alter chromatin accessibility, especially pioneer factors, while TF reporter assays reveal the transcriptional potency of TFs, particularly those activated via signaling. Each provides valuable, but partial, insight into TF activity. A combined approach is essential for a comprehensive understanding of gene regulation. While the primeTF reporter assay appears more sensitive to functionally relevant changes in many scenarios, integrating both assays will better capture the full regulatory landscape.

## METHODS

### Cell culture

mESCs (E14TG2a, #CRL-1821, ATCC), mESCs with FKBP-tagged POU5F1 (genetic background: V6.547) or SOX2 (IB10, both kindly provided by Elzo de Wit (Netherlands Cancer Institute)) (26) or mESCs with a stably integrated inducible FOXA1 overexpression cassette (23) were cultured in 2i+LIF culturing media according to the 4DN protocol (https://data.4dnucleome.org/protocols/cb03c0c6-4ba6-4bbe-9210-c430ee4fdb2c/). The reagents used were neurobasal medium (#21103-049, Gibco), DMEM-F12 medium (#11320-033, Gibco), BSA (#15260-037, Gibco), N27 (#17504-044, Gibco), B2 (#17502-048, Gibco), LIF (#ESG1107, Sigma-Aldrich), CHIR-99021 (#HY-10182; MedChemExpress) and PD0325901 (#HY-10254, MedChemExpress), monothioglycerol (#M6145-25ML, Sigma) and L-Glutamine (#25030-081, Gibco). The mNPCs used in this study were differentiated from E14TG2a mESCs and cultured in mNPC medium as mentioned previously (32).

### Prime TF reporter transfections

The prime TF reporter library was transfected as described previously (24) with some modifications. 3x10^5^ cells were seeded in a 12-well and transfected directly after plating by adding 1 μg of prime TF reporter plasmid library with 3 μl of Lipofectamine 2000 (#11668027, ThermoFisher) in 100 μl Opti-MEM (#31985070, Gibco). At least two independent biological replicates of the transfections were done on two separate days.

### TF perturbations and cell harvesting

Degradation of POU5F1 or SOX2 was induced directly after TF reporter library transfections using 500 nM dTAG-13 (#SML2601, Sigma). TFCP2L1 knockdown was performed by adding 40 nM siRNA targeting TFCP2L1 (#L-051287-01, Dharmacon) to the plasmid transfection mix during the library transfection. A non-targeting siRNA was used as a negative control (#D-001210-01, Dharmacon). FOXA1 was overexpressed by treating the FOXA1 overexpression cassette-carrying cells for 24 hours with 2 μg/ml doxycycline (#D9891, Sigma) prior to the transfection, and 24 hours after the transfection. LIF and Chiron depletion was done by plating the cells at the time of transfection in LIF or Chiron-depleted medium after washing cells twice in PBS. For the LIF depletion time-course experiments, the cells were kept in 2i+LIF medium after transfection, after which medium was changed to LIF depleted medium at 6, 3, or 1 hour before harvesting. Cells were washed twice with PBS to remove any leftover LIF. For the serum stimulation experiments, 2i+LIF medium was exchanged with medium supplemented with 20% FBS 18 hours after transfection and 6 hours before harvesting. For forskolin stimulation, 10 µM forskolin (#HY-15371, MedChemExpress) was added 18 hours after transfection and 6 hours prior to harvesting. Heat shock was induced by incubating the cells at 43 °C for 3 hours prior to harvesting the cells. In all conditions, the cells were counted and harvested 24 hours after library transfection. 50,000 cells were directly used for lysis and ATAC-seq processing, the remaining cells were resuspended for RNA isolation in 800 μl TRIsure (#BIO-38032; Bioline) and stored at -80 °C until further use.

### TF reporter barcode sequencing

RNA extraction was done using the standard procedure according to the TRIsure protocol. cDNA synthesis and barcode amplification were performed as described previously (24). The plasmid DNA was sequenced as described previously (24). The final library was sequenced on an Illumina MiSeq using 75 bp single-read sequencing yielding on average approximately 510,000 reads per sample and 789 reads per barcode.

### ATAC-seq

ATAC-seq was done using the previously established protocol (6) with slight modifications as described before (26). Quality of the libraries was checked on a Bioanalyzer before sequencing.

### ATAC-seq data preparation

ATAC-seq reads were aligned to the mm10 reference genome using BWA (v0.7.17-r1188) with the bwa mem -M command (63). Alignments were filtered using Samtools (v1.9) with samtools view -h -b -q 10 to retain high-quality mappings (64). Duplicate reads were removed using the MarkDuplicates function in Picard Tools (v2.12.0) with the argument REMOVE_DUPLICATES=true. Peaks were called using the callpeak function using macs2 (v2.2.6) (65).

### Sox2 accessibility tornado plots

Tornado plots were generated using deeptools (v3.5.1) (66). Bigwig files were created by normalizing computing size factors for each sample using the estimateSizeFactors function of DESeq2 (v1.30.1) (67). To select the 1000 best-matching SOX2 binding sites in the genome, the peaks were scanned with the SOX2 motif (M06121_1.94d, CIS-BP (build 1.94)) using FIMO (68), and then the sites with the 1000 smallest p-values were selected.

### chromVAR analysis

chromVAR (v1.12.0) was run using peaks with lengths set to 500 bp using the slop function from bedtools (v2.26.0) (69). GC bias was added using the addGCBias function from chromVAR, and peaks were filtered using the filterPeaks function from chromVAR, setting non_overlapping to true. Peak count data was overlapped with selected motif position weight matrices (PWMs) as described in Trauernicht *et al.* in Table S1 (23) using the matchMotifs function from the motifmatchr package (v1.12.0). For prime TF reporters based on published sequences (i.e., non-synthetic reporters without clearly selected TFBSs), we selected PWMs based on Lambert *et al.* (70) as selected for Figure 2A in that publication. These PWMs were also used to calculate chromVAR deviation scores for all non-selected TFs in the mNPC differentiation experiment. Background peaks were identified using 2000 iterations using the getBackgroundPeaks function from chromVAR, as recommended before (25). Deviations were calculated using the computeDeviations function and extracted using the deviations function from chromVAR. For downstream analyses, deviation scores were averaged across replicates. To identify statistically significant differential accessibility between tested conditions we used a previously optimized workflow that combined chromVAR with limma (v3.46.0) (25) using the eBayes function.

### TF reporter activity computation and primetime analysis

TF reporter activities of individual replicates were calculated as described previously (12). Briefly, barcode counts were normalized by read depth by computing reads per million and then normalized to the plasmid DNA counts. To identify statistically significant up- and downregulated TF reporters, *primetime* (v0.4) was run using default parameters and executed as described previously (24). Samples were scanned and filtered using *primetime*’s quality check metrics, ensuring low plasmid DNA bleedthrough, high barcode correlation and high biological replicate correlation.

## DATA AVAILABILITY

Raw ATAC-seq data and the reporter assay sequencing data is available through SRA under BioProject ID PRJNA1357637. Laboratory notes and supplementary raw data are available at Zenodo: https://doi.org/10.5281/zenodo.18020009. Original code and analysis pipelines are available at GitHub: https://github.com/mtrauernicht/ATAC_TF_reporter_comparison. A released version of the code is available at Zenodo: https://doi.org/10.5281/zenodo.18019661.

## AUTHOR CONTRIBUTIONS

Max Trauernicht: Experiments; Data analysis; Manuscript writing. Vinícius H. Franceschini-Santos: Data analysis, Manuscript review. Teodora Filipovska: Experimental setup, Manuscript review; Bas van Steensel: Manuscript writing; Project supervision.

## ACKNOWLEDGMENTS

We thank members of our laboratories for helpful comments and the NKI Genomics and Research High-Performance Computing core facilities for technical support. We thank the lab of Elzo de Wit for Sox2 and Oct4 degron cell lines. We also thank Hans Teunissen (Netherlands Cancer Institute) for support during the execution of the ATAC-seq protocol, and Teun van den Brand (Netherlands Cancer Institute) for supporting the computational analyses. This work was funded by the Oncode Institute and the European Union (European Research Council Advanced Grant RE_LOCATE, 101054449). Views and opinions expressed are however those of the author(s) only and do not necessarily reflect those of the European Union or the European Research Council. Neither the European Union nor the granting authority can be held responsible for them. Research at the Netherlands Cancer Institute is supported by an institutional grant of the Dutch Cancer Society and of the Dutch Ministry of Health, Welfare and Sport. The Oncode Institute is partially funded by the Dutch Cancer Society.

## EXPANDED VIEW FIGURES

**Figure EV1:**
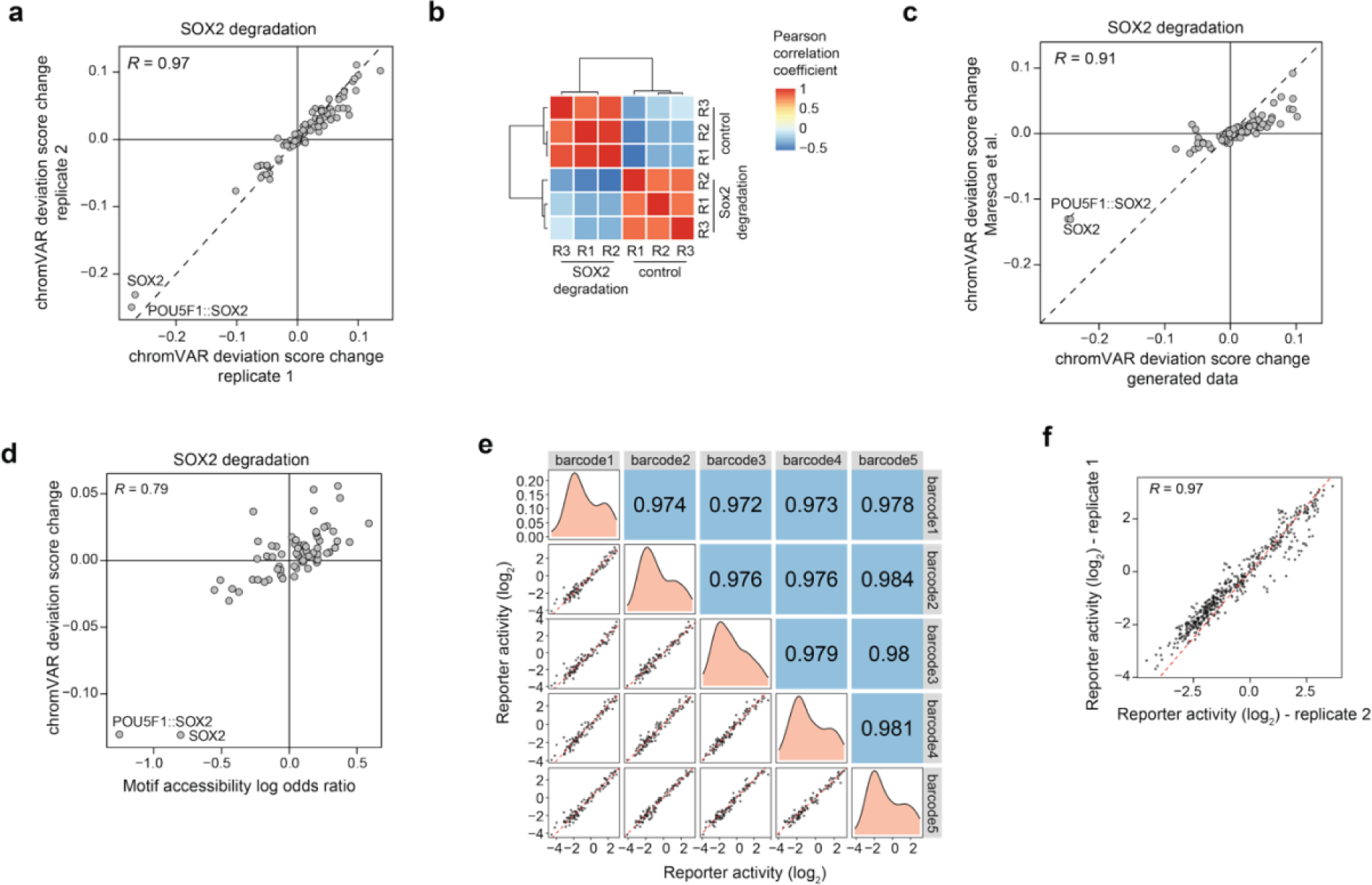
Quality control of chromVAR and primetime. a) Correlation of chromVAR deviation score changes upon Sox2 degradation between two biological replicates. b) Heatmap showing Pearson correlation coefficients of chromVAR deviation scores between replicates for Sox2 degradation and control conditions. c) Correlation of chromVAR deviation score changes upon Sox2 degradation between previously published data and data generated in this study. d) Correlation between chromVAR deviation score changes and motif accessibility log odds ratio from previously published data. e) Correlation of reporter activity between the five barcodes used in the prime TF reporter library. f) Correlation of reporter activity upon Sox2 degradation between two biological replicates.

**Figure EV2:**
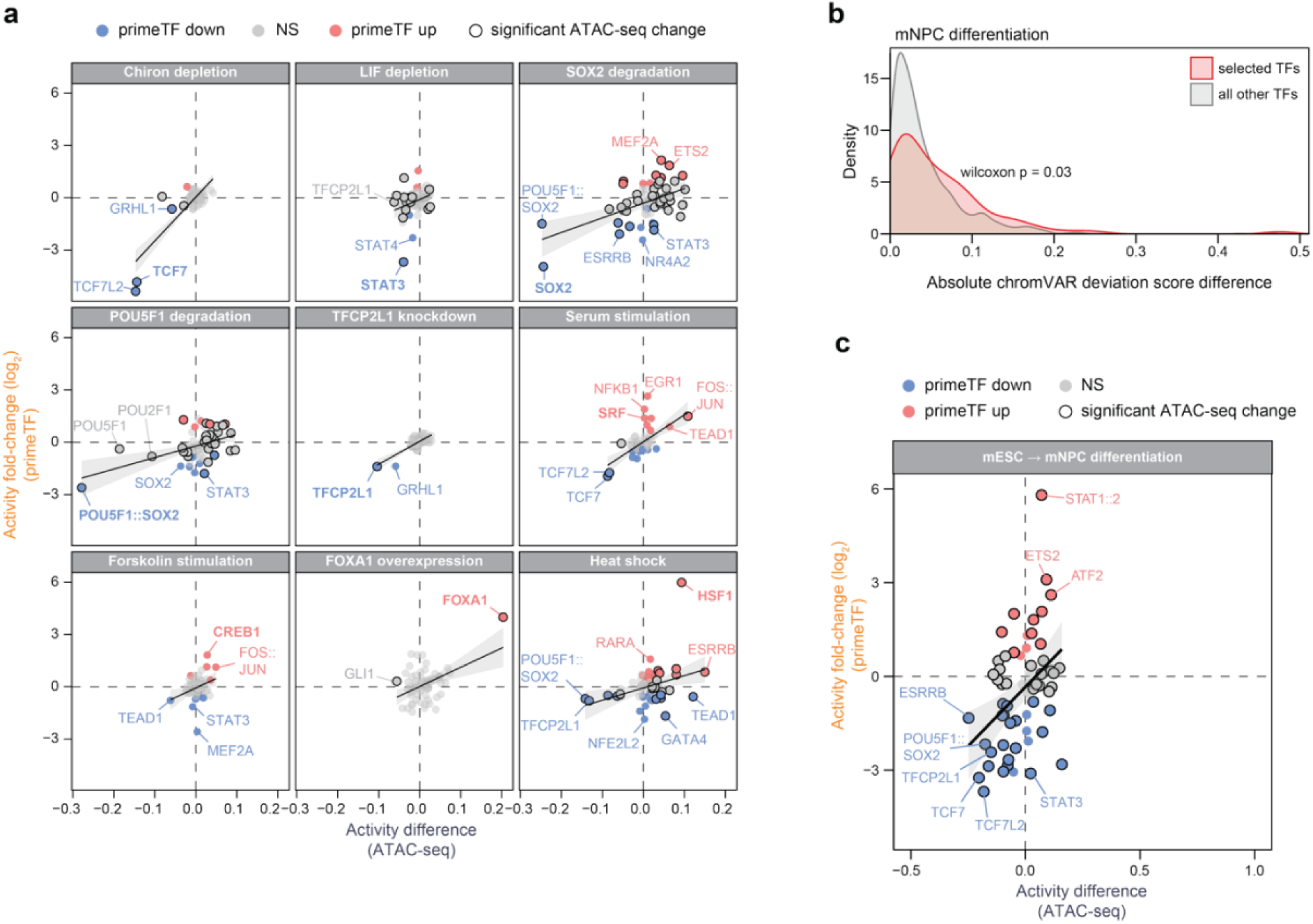
Extended analyses using only prime reporters. a) Same correlation plot as depicted in Figure 4a, filtering for only TFs with prime reporters. b) Density plot of the absolute chromVAR deviation score differences for the 100 selected TFs (red) and all non-selected TFs (grey). Statistical significance between these two groups was tested using Wilcoxon rank-sum test. c) Same correlation plot as depicted in Figure 5c, filtering for only TFs with prime reporters.

